# Early maturation of sound duration processing in the infant brain

**DOI:** 10.1101/2023.01.24.525349

**Authors:** Silvia Polver, Brigitta Tóth, Gábor P. Háden, Hermann Bulf, István Winkler

## Abstract

The ability to process sound duration is crucial already at a very early age for laying the foundation for the main functions of auditory perception, such as object perception and music and language acquisition. With the availability of age-appropriate structural anatomical templates, we can reconstruct EEG source activity with much-improved reliability. The current study capitalized on this possibility by reconstructing the sources of event-related potential (ERP) waveforms sensitive to sound duration in four- and nine-month-old infants. Infants were presented with short (200 ms) and long (300 ms) sounds equiprobably delivered in random order. Two temporally separate ERP waveforms were found to be modulated by sound duration. Generators of these waveforms were mainly located in primary and secondary auditory area and other language-related regions, such as the superior temporal and the inferior frontal gyri. The results show marked developmental changes between four and nine months, partly reflected by scalp-recorded ERPs, but appearing in the underlying generators in a far more nuanced way. The results also confirm the feasibility of the application of anatomical templates in developmental populations.

## 1. Introduction

One of the main characteristics of auditory information is that sound unfolds in time. Therefore, the integration of information across time is crucial for perception on multiple timescales ^[1]^. Infants are sensitive to several temporal sound features at birth, such as gaps, sound onset and offset, and changes in sound duration ^[2–4]^. Therefore, these abilities are thought to be innate and to support the foundation for language acquisition ^[2,5,6]^. Similarly to adults, infants also utilize temporal cues, such as phoneme duration for segmenting sound sequences ^[7]^, including finding words within continuous speech^5^. In this sense, the ability to process rapidly changing sounds has been suggested to be a critical skill for language development ^[8,9]^. However, despite their importance, the mechanisms underlying the processing of auditory temporal features are still poorly understood. The present study aimed to investigate the early development of the processing of an important auditory temporal feature: sound duration. To this end, auditory event-related brain potentials (ERP) and their neural generators elicited by tones and noises of different duration were compared between four and nine-month-old infants.

ERPs have been long used to study the neural substrate of auditory temporal processing and their development in infancy ^[2,3,10]^. Specifically, some previous ERP studies have shown that the newborn brain is sensitive to sound duration as well as to detect unexpected changes in the duration of a repeating sound ^[3,11,12]^. The adult ERP components sensitive to sound duration comprise the P1-N1-P2-N2 complex ^[3]^. The adult P1–N1–P2–N2 sequence begins at around 40-50 ms after stimulus onset and lasts until ca. 150-250 ms ^[13]^. Similar waveforms have been observed in newborns, infants, and in 5 to 10 years old children [2]. However, it is debated whether the underlying functional mechanisms and brain areas involved are the same as those observed in adults ^[14]^ .

Generally, neonatal ERPs exhibit a large positive waveform at midline starting at about 100 ms and ending 450 ms from stimulus onset, followed by a low-amplitude negative deflection at approximately 450-600 ms (N450). These waveforms are thought to reflect precursors of the P2 and N2 ^[15]^. At about three months of age, the P2 is reportedly split by the appearance of a negative deflection, and between three and six months, another deflection peaking at about 250-280 ms (N250) appears. The two negative waveforms are assumed to represent precursors of the adult N1 and N2, respectively ^[15]^. By six- to nine months of age the separation between the N1 and the N2 precursor waveforms becomes well-defined, making the infantile ERPs’ morphology resemble the P1-N1-P2-N2 complex ^[14,16,17]^. The developmental trajectories of ERP waveforms point toward the presence of underlying developmental milestones from three to nine months of age ^[14,16,17]^. Unfortunately, studies directly addressing changes along development are still scarce.

However, scalp recordings do not directly reflect the underlying generators. Sound perception activates multiple pathways in the brain. Thus scalp-recorded waveforms may represent summed activity from several generators ^[18,19]^. Moreover, different ERP generators may have different maturational courses ^[15]^. Therefore, discerning and locating these generators and addressing their developmental trajectory provides important information for anchoring functional changes in brain maturation and plasticity. In adults, generators of the P1-N1-P2-N2 complex have been found to reside in the primary auditory cortex (Heschl’s gyrus), the superior temporal gyrus (STG), in thalamo-reticular systems, in the planum temporale (encompassing Wernicke’s region), and in supratemporal auditory cortices ^[18,20–22]^. Regarding developmental populations, source analysis revealed that presenting syllable to 6-months-old infants activated bilaterally the auditory and frontal cortices, and the anterior cingulate cortex ^[23]^. 6-months-old infants also show activity in supratemporal and frontal areas in response to sound discrimination ^[24]^. However, until recently, the lack of age-appropriate templates has complicated the application of source localization in infants, because the quality of age-appropriate head models determines the accuracy of source localization ^[19]^. Here we rely on the recently released infant templates ^[25]^ derived from the Neurodevelopmental MRI Database ^[26]^ and made ready for use by their implementation in the Brainstorm software ^[27]^.

We investigated duration-sensitive auditory ERP waveforms and their sources to establish their developmental changes between four and nine months of age. We chose these two time points, because they bracket important milestones in the development in ERP morphology (see above). Infants were presented with four sounds differing in duration (200 vs. 300 ms) and sound quality (harmonic tones vs. white noise). We expected to find significant changes in ERP morphology between the two age-groups in accordance with previous descriptions of ERP development. Indeed, in contrast to the simpler ERP component structure at four months, by nine months of age we expected to find precursors of the adult ERP complex in the form of a positive deflection, accompanied by two negative ones coinciding with the N250 and the N450 latencies ^[15]^. We then asked whether either or both waveforms show sensitivity to sound duration and whether sensitivity to sound duration is dependent on the quality of the sound (harmonic tones vs. noise segments).

To our knowledge, this is the first study applying age appropriate templates published by O’Reilly and colleagues (2020) ^[25]^ to reconstruct the sources of auditory ERPs in infants. We expected to find the main sources of duration-sensitive activity in brain areas belonging to the auditory brain network, including primary auditory cortex (PAC), and activity in network areas previously shown to be sensitive to sound duration, such as the STG and inferior frontal areas ^[28]^. Further, in line with our expectations for scalp-recorded ERPs, we hypothesized that nine-months-old infants show stronger activity in a more distributed network comprising fronto-central areas compared to four-months-old infants in the time window(s) of the effect(s) found over the scalp.

## 2. Results

Figure 1 shows the grand average ERP difference responses and their neural generators for tones (panels A, B, and C), while Figure 2 shows the same for noise segments (panels A, B, and C) at four and nine-month of age.

**Figure 1:**
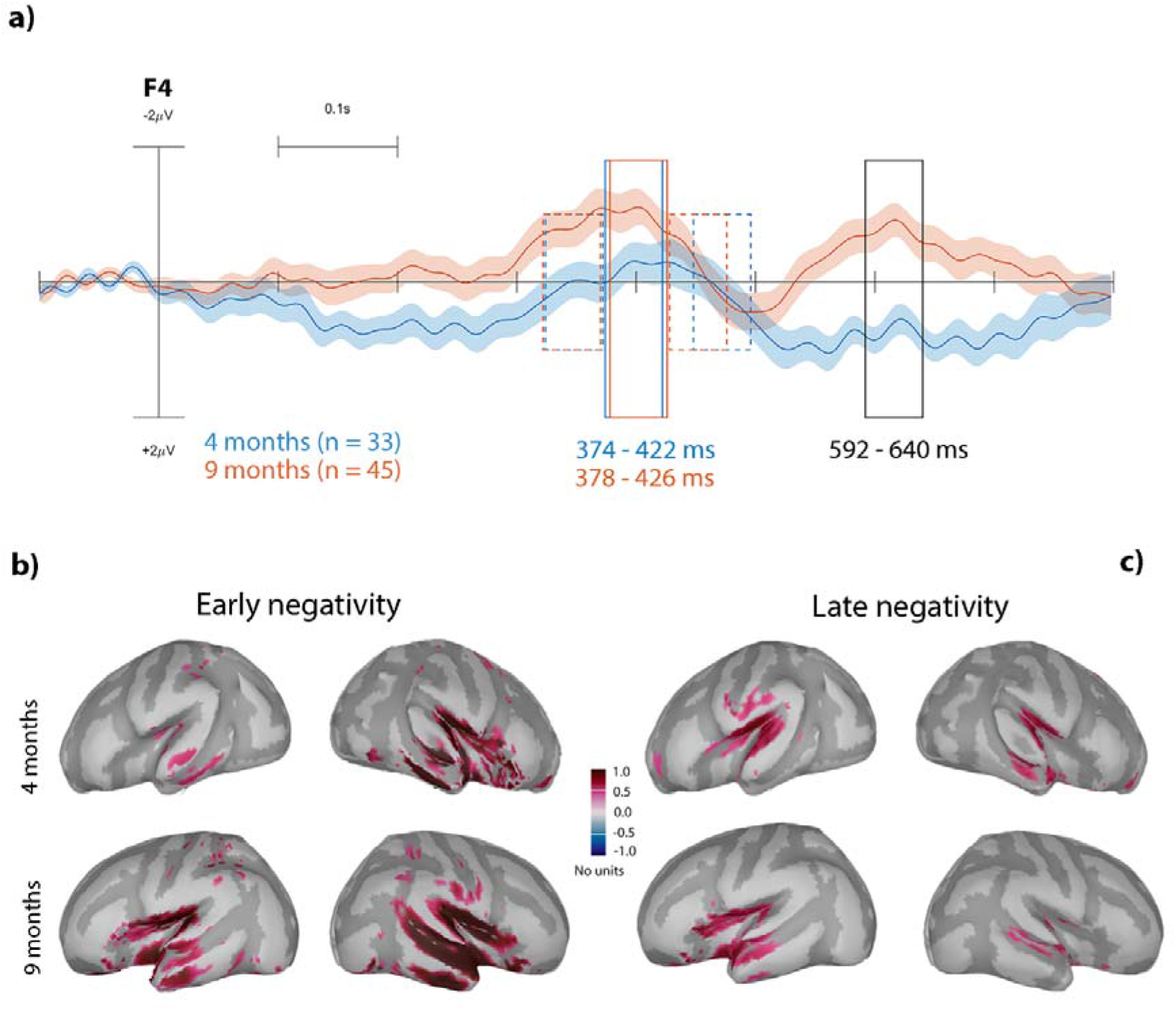
Grand average ERP difference waveforms elicited by tones. Panel **A)** overplots the right frontal (F4) ERP responses of the two age groups (marked by line colors). The early and the late measurement windows are shown with rectangles drawn with solid lines (the exact range shown under the rectangle). The rectangles with dashed lines refer to the reference interval used for calculating the amplitudes for the early waveform. Panels **B)** and **C)** show the distribution of the average current source density (CSD) over the inflated cortex during the early (panel **B**) and late (panel **C**) duration-sensitive ERPs elicited by tones. The upper row presents the four-months old infants’ data, the lower that of the nine-month-olds. Color scale is at the center of the cortical distribution panels.

**Figure 2:**
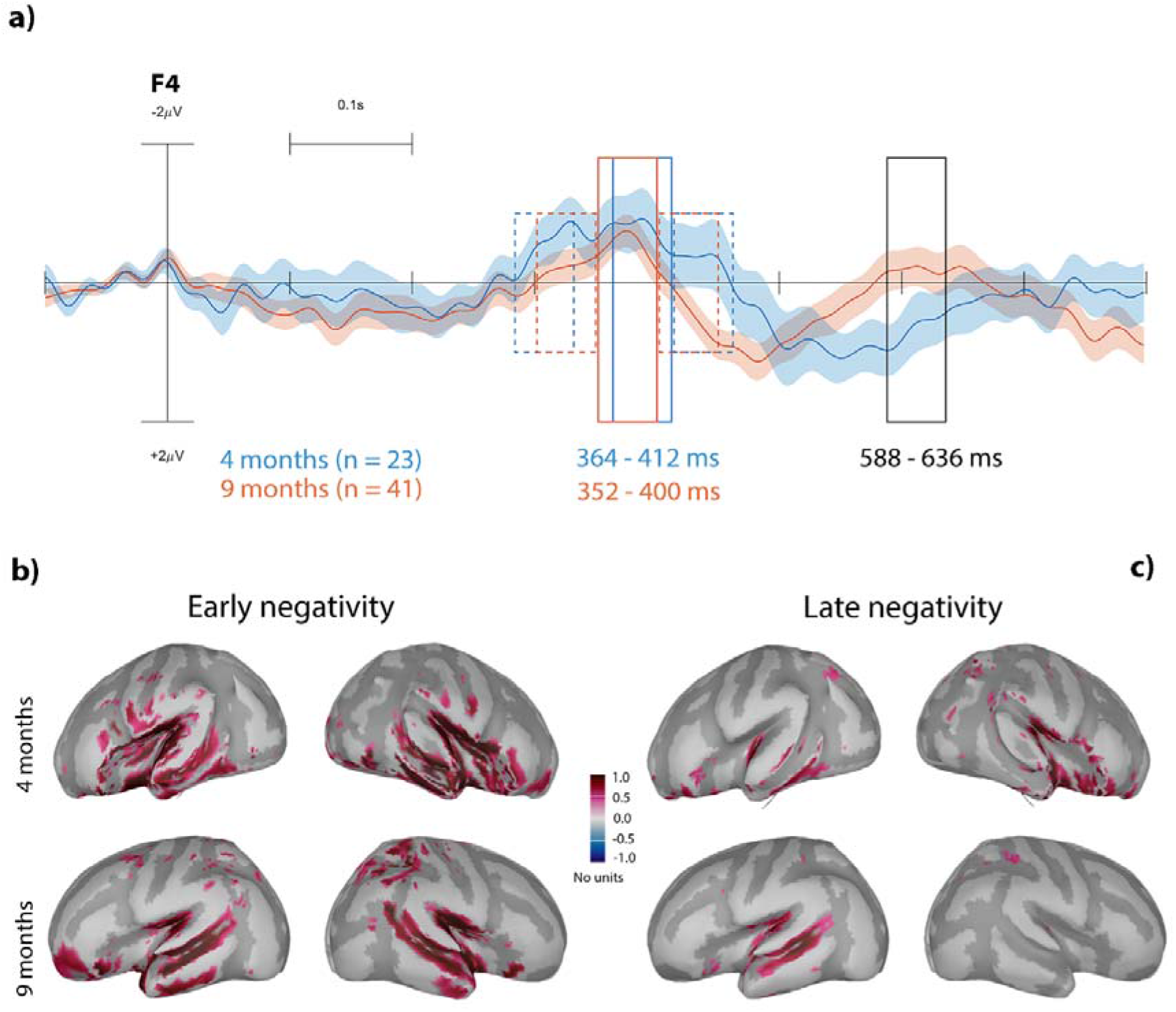
Grand average ERP difference waveforms elicited by noise segments. Panel **A)** overplots the right frontal (F4) ERP responses of the two age groups (marked by line colors). The early and the late measurement windows are shown with rectangles drawn with solid lines (the exact range shown under the rectangle). The rectangles with dashed lines refer to the reference interval used for calculating the amplitudes for the early waveform. Panels **B)** and **C)** show the distribution of the average current source density (CSD) over the inflated cortex during the early (panel **B**) and late (panel **C**) duration-sensitive ERPs elicited by noise. The upper row presents the four-months old infants’ data, the lower that of the nine-month-olds. Color scale is at the center of the cortical distribution panels.

### 2.1. Early negative difference

For tones, the amplitude measurement window for the peak of the early negativity was set from 374 to 422 ms at four months and from 378 to 426 ms at nine months (Figure 1A). Results of the ANOVA showed significant main effects of ANTERIOR-POSTERIOR (F(2,684) = 28.89, p < 0.0001, ηp^2^ = 0.08) and AGE GROUP (F(1,684) = 12.9, p = 0.0008, ηp^2^ = 0.02), and a significant interaction between ANTERIOR-POSTERIOR and AGE GROUP (F(2,684) = 8.14, p = 0.0008, ηp^2^ = 0.02). The mean amplitude of the four-months-old responses was lower than that of the nine-months-olds (AGE GROUP main effect): −0.08 μV vs. −0.28 μV, respectively. Post-hoc Tukey’s pairwise comparisons for the significant ANTERIOR-POSTERIOR main effect revealed that frontal (t(684) = −5.82, p < 0.0001) and central (t(684) = −6.45, p < 0.0001) sites are characterized by significantly higher early negative responses compared to posterior sites (−0.306 μV and −0.386 μV vs. 0.09 μV, respectively). Post-hoc ANOVAs conducted for the significant ANTERIOR-POSTERIOR × AGE GROUP interaction revealed that the groups differed only at the central line of electrodes (F(1,232) = 36.71, p < 0.0001, ηp^2^ = 0.14) with four-months-old infants producing significantly lower-amplitude (less negative) responses than nine-months-olds (−0.08 μV vs. - 0.6 μV).

For noise segments, the amplitude measurement window for the peak of the early negativity was set from 362 to 412 ms at four months and from 352 to 400 ms at nine months (Figure 2A). Results of the ANOVA revealed only a significant ANTERIOR-POSTERIOR main effect (F(2,558) = 76.82, p < 0.0001, ηp^2^ = 0.22). Post-hoc pairwise comparisons revealed that frontal (t(558) = −10.48, p < 0.0001) and central (t(558) = −9.71, p < 0.0001) sites are characterized by significantly higher early negativity responses compared to posterior sites (−0.44 μV and −0.38 μV vs. 0.39 μV, respectively).

#### Generators of the early negative difference

Distributions of the grand-average current source density (CSD) differences are shown on Figure 1B and Figure 2B, for tones and noise segments, respectively. Significant AGE GROUP effects were found for tones in the right temporal cortices (right MTG, STG, and the bank of the STS) and in the IFG (left pars opercularis). Significant AGE GROUP effects were found for noise segments in all left partitions of the IFG (left pars opercularis, triangularis, orbitalis) and in the right bank of the STS. All significant statistical findings are listed in Table 1. of the Supplementary Materials.

**Table 1.** Mean number of epochs remaining after preprocessing reported separately for STIMULUS DURATION, STIMULUS TYPE, and AGE GROUP.

|  | Short tones | Long tones | Short noises | Long noises |
| --- | --- | --- | --- | --- |
| four-months-old | 133.53 (SD = 15.6) | 133.96 (SD = 15.13) | 131.04 (SD = 19.69) | 131.78 (SD = 18.16) |
| nine-months-old | 134.82 (SD = 21.47) | 134.17 (SD = 18.95) | 135.04 (SD = 30.24) | 136.87 (SD = 30.08) |

### 2.2. Late negative difference

For tones, the amplitude measurement window for the late negativity was set from 592 to 640 ms (Figure 1A). Results of the ANOVA showed a significant main effect of AGE GROUP (F(1,684) = 11.41, p = 0.002, ηp^2^ = 0.02), and a significant interaction between the ANTERIOR-POSTERIOR and AGE GROUP (F(2,684) = 11.09, p = 0.0001, ηp^2^ = 0.03). The mean amplitude of the four-months-old responses was positive in contrast to that of the nine-months-olds (AGE GROUP main effect): 0.3 μV vs. −0.18 μV. The post-hoc ANOVA on the ANTERIOR-POSTERIOR and AGE GROUP interaction revealed that the groups significantly differed only at frontal (F(1,232) = 20.6, p < 0.0001, ηp^2^ = 0.08) and central electrodes (F(1,232) = 13.59, p = 0.0002, ηp^2^ = 0.06), with the four-months-old infants responses characterized by positive mean amplitudes (0.55 μV and 0.51 μV for frontal and central sites, respectively) compared to that of the nine-months-olds (−0.52 μV and −0.33 μV for frontal and central sites, respectively).

For noise segments, the amplitude measurement window for the late negativity was set from 588 to 636 ms (Figure 2A). Results of the ANOVA revealed a significant interaction between ANTERIOR-POSTERIOR and AGE GROUP (F(2,558) = 17.09, p < 0.0001, ηp^2^ = 0.06). The post-hoc ANOVAs on this interaction revealed that the groups differed at all three electrode lines. At frontal (F(1,190) = 8.37, p = 0.004, ηp^2^ = 0.04) and central (F(1,190) = 10.3, p = 0.001, ηp^2^ = 0.05) electrodes, four-months-old infants (0.44 μV and 0.18 μV respectively) produced positive responses as opposed to nine-months-olds (−0.38 μV and −0.6 μV respectively). In contrast, four-months-old infants were characterized by negative responses in the late negativity while nine-months-olds by positive ones (−0.65 μV vs. 0.48 μV) at posterior sites (F(1,190) = 16.15, p = 0.0008, ηp^2^ = 0.08).

#### Generators of the late negative difference

Distributions of the grand-average current source density (CSD) differences are shown on Figure 1C and 2C. The effect of AGE GROUP on the late negative difference to tones was significant in temporal areas (bilateral STG, and right PAC), in the right parietal SMG region, and in the left pars triangularis of the IFG. Significant effects of AGE GROUP on the late negative difference to noise segments were observed in temporal areas (bilateral STG, and PAC, left bank of the STS, ITG, MTG), in the right SMG, in the left lateral OFG, in the right medial OFG, and in the left pars orbitalis of the IFG. All significant statistical findings are listed in Table 2. of the Supplementary Materials.

## 3. Discussion

Results of the current study showed that the brain of four- and nine-months-old infants is sensitive to sound duration, as reflected by ERPs elicited by short, isolated sounds. Effects of maturation were found for the scalp-recorded waveforms and the corresponding source activity. Both scalp-recorded ERPs and the related source activity became stronger and spatially better defined between four and nine months of age. Generators of the event-related potentials sensitive to sound duration were located in the ventral frontotemporal auditory pathway. Details of the results are discussed below.

Two successive fronto-centrally negative sound-duration sensitive waveforms were identified in the ERPs elicited by nine-months old infants. In contrast, only the first of these two negativities were present in the responses of four-months-olds with similar, 350-400 ms peak latencies in the two age groups. The late negativity peaked between 580 and 650 ms in the nine-months-olds over fronto-central scalp sites. The emergence of a second sound duration sensitive waveform between four and nine months of age may reflect that this information is further processed in the latter age group. The ERP results are generally congruent with reports of waveforms related to the N250 and N450 in infant studies with duration manipulation ^[29,30,3,15]^. While we did observe slightly longer latencies in the present study compared to some of the previous ones, this may be due to variability in the maturational stages of auditory brain networks ^[31]^. The fronto-centrally maximal scalp distributions of the current waveforms are also compatible with those reported in previous literature ^[31,32,3]^. This fronto-central gradient has been suggested to be part of the developmental trajectory of auditory ERPs ^[6]^.

In contrast to some other studies, we did not observe lateralized effects for the early negativity. Different generators underlying the same ERP can have different maturational pathways ^[31]^. This may explain the mixed reports of age effects on lateralization ^[33,24]^. Thus, the contrasting age-related differences in generator activity may be related to differential maturational trajectories ^[35]^. This hypothesis is strengthened by previous reports highlighting that ERP components underlying duration perception are not unitary in nature and reflect dynamic processes involving multiple brain regions ^[36]^.

The source-localized results show that the generators underlying the early negativity are distributed exclusively within the ventral auditory pathway consisting of bilateral temporal (PAC, MTG, STG, ITG) and inferior frontal cortices (see Figure 1B and 2B). The neural generators of the late negativity were located in overlapping areas (see. Figure 1C and 2C). However, they showed more focal distribution within primary and secondary auditory areas. The duration sensitive sources found in this study are consistent with those reported in the literature in response to both speech sounds and tones: the superior temporal, supramarginal, and inferior frontal gyri, and primary and secondary auditory cortical areas [31,37,38,21,34,39]. This points towards the reliability of the use of anatomical templates in developmental samples, which represents an important turning point in studying electrical brain activity in infants.

A somewhat surprising result was the similarity between the scalp-recorded ERP morphologies of four- and nine-months-old infants. Because the infant brain is subject to dramatic development during this period ^[31]^, one may expect the morphology of all ERP responses to change accordingly. However, similarity between the results in the two age groups sheds light on the central role the processing temporal information plays in sound processing: the processing of temporal features is more consistent than that of spectral features throughout development and even its neural substrate may not change dramatically from infancy to adulthood. Thus, whereas pitch discrimination initially requires very large separation between the sounds to elicit reliable discriminative brain activity ^[40]^, temporal features are reliably detected already at birth ^[4,10]^ and responses sensitive to different sound durations are quite similar between neonates and adults ^[3]^.

The underlying generators show a more nuanced picture. As seen in Figure 1 and 2, the cortical distribution of the sound duration sensitive generators active during the early scalp-recorded negativity are almost identical in the two age groups. The exception is represented by the involvement of inferior frontal regions, suggesting the presence of brain maturational mechanisms. Supporting this assumption, we found statistically significant increments in current source density within the time window of the early negativity: increased activity was observed in the right hemispheric part of the previously described fronto-temporal network (see Supplementary Table 1). These mechanisms may be based on either higher efficiency of the primary and secondary auditory sensory cortices (temporal brain regions), and/or the increased involvement of additional regions such as the inferior frontal cortex.

The cortical distribution of the neural activity within the late negativity time window also strongly overlapped between age groups. However, substantial age-related differences were found in left primary and secondary auditory areas (bilateral STG, bilateral PAC, left bank of the STS, left ITG, and left MTG). In addition, we observed age related differences in the associative right SMG and in the OFG (Supplementary Table 2). These distributions indicate that maturation aids auditory network development in a quantitative manner at the later stages of duration processing.

Age-related effects on duration-sensitive ERPs and their generators may stem from multiple types of maturational processes ^[15]^. For example, myelination of auditory pathways plays a crucial role by enabling rapid coordination among neurons ^[41]^. The timeline of myelination in auditory pathways mirrors key developmental milestones in auditory perception such as the appearance of the P1-N1-P2-N2 ^[41]^. Regarding the involvement of frontal regions, despite the widely accepted notion that prefrontal areas show prolonged development compared to sensory cortical regions ^[35]^, some studies show that already at birth infants recruit frontal areas during processing speech sound ^[42,43,44]^. Moreover, temporal and frontal regions do not develop independently but show high correlations in their developmental course ^[45]^. These findings suggest that the frontal cortices and their connections to auditory sensory regions (temporo-parietal areas) play a major role in language acquisition. In infants, frontal and temporo-parietal regions are already well connected and these connections are hypothesized to underlie the early processing of speech [46]. Both bottom-up and top-down connections are necessary for language ^[47]^. We hypothesize that the first duration-sensitive component to reflect a first prodromal involvement of frontal cortices in auditory temporal processing (also related to language acquisition), the late component may then reflect feedback from frontal cortices to temporal areas. This hypothesis is supported by the involvement of primary auditory cortices in the generation of the later duration-sensitive component. In this sense, the late component might be reflecting a maturational step of duration processing which might not yet be stable at four months.

The current results provide new information on the development of the processing of an important temporal parameter, sound duration. The ERP results are generally compatible with those from previous studies. The more reliable source reconstruction offered by the recently released infant structural anatomical templates ^[25]^ provided new insights into the generators underlying duration processing and their development, allowing one to speculate about the functional and developmental significance of this neural activity. Naturally, these initial hypotheses require further studies for confirmation and specification.

## 4. Methods

### 4.1. Participants

Infants were recruited as part of a broader longitudinal study aimed to assess the role of infant-directed speech in speech acquisition. In this manuscript, we will refer only to two separate age cohorts: the first one comprised 64 (20 males) four-months-old, the second cohort 63 (19 males) nine-months-old healthy full-term infants. All infants were firstborns, none of them twins. The average age at the recording time was 4.22 months (SD = 0.2) for the four-months-old and 9.52 for the nine-months-old group (SD = 0.36). Data recorded from 31 four-months-old infants and from 18 nine-months-old infants was excluded due to the presence of excessive artifacts (see Sections 4.3 and 4.4). Thus, 33 four-months-old and 45 nine-months-old infants’ data were included in the final sample. Due to the longitudinal nature of the study, 9 of infants in the final sample were tested at both four and nine months.

Recordings were carried out at the sound attenuated infant laboratory of the Institute of Cognitive Neuroscience and Psychology, Research Centre for Natural Sciences, Budapest, Hungary. Informed consent was obtained from one (mother) or both parents. The study was conducted in full accordance with the Declaration of Helsinki and all applicable national and international laws, and it was approved by the United Ethical Review Committee for Research in Psychology (EPKEB), Hungary.

### 4.2. Stimuli and procedure

All infants were presented with shorter (200 ms) or longer (300 ms) sounds grouped into two separate blocks each consisting of 150 short and 150 long sounds delivered in random order. One of the stimulus blocks presented tones of 500 Hz base frequency with 3 harmonics of 50, 33, and 25 percent amplitude, respectively, summed linearly together (‘tone’ condition). The other stimulus block comprised frozen white noise segments (‘noise’ condition). All sounds were presented at approximately 70 dB SPL loudness and the sounds were attenuated by 5 ms long raised-cosine onset and offset ramps. The onset-to-onset interval was 800 ms. The tone condition was always delivered first, and there was a short rest of 30-60 seconds between conditions. Stimuli were created and delivered using Matlab (MathWorks Inc., Natick, MA, USA) and Psychtoolbox ^[48]^ softwares. The sound signal from the computer was amplified by a Yamaha A-S301 amplifier (Hamamatsu, Japan) and presented through a pair of speakers (Boston Acoustics A25, Woburn, MA, USA). The speakers were positioned ca. 1.75 meters in front of the participant, 70 cm apart from each other.

The infant sat comfortably in her mother’s lap while the experimenter employed toys to keep the infant facing toward the loudspeakers and her attention away from the electrode net. The mother was listening to music through closed can audiometric headphones to isolate her from the experimental stimulation. If the infant became fussy, the playback was stopped, and the experimenter attempted to pacify her. If pacifying was successful the playback restarted from the beginning, otherwise, the experiment was discontinued. Because some infants did not complete both stimulus blocks, 33 four-month-old and 45 nine-month-old infants’ data was analyzed for the tone condition, and 23 four-months-old and 41 nine-months-old infants’ data for the noise condition.

The experiment reported here was presented first within a session combining multiple experiments. Data from the other experiments and the associations across results will be published separately after collecting the complete data set. As the current experiment was always the first one presented, the other experiments are not listed.

### 4.3. EEG data acquisition and preprocessing

EEG was recorded with a 60-channel HydroCel GSN net (64 channel v1.0 layout, geodesic; channels 61-64 were connected to the ground in the small pediatric caps used) and a GES 400 DC amplifier passing the digitized signal to a computer running the NETSTATION v5.4.1.1 software (both Electrical Geodesics, Eugene, OR, USA). Signals were recorded online at a sampling rate of 500 Hz with the Cz reference (DC, no on-line filtering). Electrode impedance during recording was kept below 50 kΩ.

The recorded signals were imported to MATLAB (Mathworks, Natick, MA; ver. 2021a) and processed using EEGLAB ^[49]^. First, data were band-pass filtered between 0.1 Hz and 30 Hz using a finite impulse response (FIR) filter (Kaiser windowed, Kaiser β=5.65, filter length 4530 points). Data were then cleaned using the Artifact Subspace Reconstruction (ASR) algorithms in EEGLAB ^[50]^: electrodes were removed and stored for subsequent interpolation if either they were not functioning for longer than 5 seconds, or if their correlation with neighboring electrodes was under 0.8. Correlation was computed for each electrode with the RANdom SAmple Consensus (RANSAC) reconstruction. The reason behind the latter choice was the assumption that nearby channels should be characterized by fairly similar activity. Signals from the remaining electrodes were then entered into a Principal Component Analysis (PCA) with a 5-s sliding window, classifying PCs into high-variance (10 SD from clean data) or normal-variance sets. High-variance parts of the data, which correspond to artifacts, were then reconstructed based on normal-variance subspace components. Infants’ data were rejected either if more than 9 channels were removed or if more than 2 removed channels were neighbors. This led to the exclusion of 28 four-months-old and 14 nine-months-old infants’ data from the analysis. To further remove artifacts, an Independent Component Analysis (ICA) was performed on 45 PCA-reduced components after re-referencing them to the average reference. The topographical distribution and frequency contents of the ICA components were visually inspected, and the components constituting oculomotor, cardiac, muscular, or line artifacts were manually removed. The maximum number of rejected components was set at 11. After these steps, electrodes removed in the first artifact rejection step were spherically interpolated with the help of the complete electrode matrix.

### 4.4. ERP analysis

From the continuous EEG records, epochs were extracted between −100 and 800 ms relative to the sound onset, separately for the two STIMULUS DURATIONs (short- and long-duration sounds), STIMULUS TYPEs (tones and noises), and AGE GROUPs (four- and nine-month-olds). The average voltage in the −100 and 0 ms window served as the baseline. A threshold of +/− 150 μV was set to reject epochs with abrupt amplitude changes, while thresholds of 100 μV over the whole epoch for slope and 0.5 r^2^ were set to remove epochs with step-like artifacts and linear trends. Infants’ data was not used if the remaining number of epochs for each STIMULUS DURATION was under 75 for either STIMULUS TYPE. This led to the additional exclusion of 3 four-months-old and 4 nine-months-old infants’ data. The final sample thus consisted of 33 four-months-old infants, of which 10 did not complete the noise condition, and 45 nine-months-old infants, of which 4 did not complete the noise condition. Epochs of each remaining infant were averaged separately for each STIMULUS DURATION, STIMULUS TYPE, and AGE GROUP. The mean number of epochs are reported in Table 1. No significant differences were found between average epoch numbers of conditions and group.

For ERP analysis, we first calculated difference waveforms by subtracting the average ERP elicited by short stimuli from the average ERP elicited by the corresponding long ones. In accordance with previous findings ^[29,30,3,15]^, we observed two successive negative peaks in the difference waveforms (Figure 1 and 2; early and late negativity: N250 and N450, respectively in Kushnerenko et al., 2002 ^[15]^). For component latencies, the right frontal (F4) grand average (GA) waveform was visually inspected. F4 was selected as the signal from this electrode typically shows the highest difference amplitude^15^.

Looking at Figure 1A and at Figure 2A the early negative difference peak elicited by four-month-olds barely passes the baseline. However, judging from the voltages surrounding the peak, the negative waveform may be riding on a positive shift or a wide positive waveform. Therefore, we decided to measure the early component’s amplitude by subtracting from the average voltage measured from a 48 ms long window centered on the peak (the most negative point in the GA response at F4) the average voltage in two windows of 48 ms duration, one on each side of the peak, separately for each stimulus type and age group. The latencies of the two baseline side-windows were determined by starting from adjacent positions on both sides of the peak, then separately sliding the windows in 24 ms steps away from the peak until their average voltage in both side-window was significantly lower (Student’s t-test; p < 0.01) than that of the peak window. After establishing the windows (separately for the two stimulus-types and two age groups; see Figure 1A and Figure 2A), the early peak’s amplitude was measured from the same windows in each infant’s individual average.

Amplitude measurement windows of the late negative waveform were established from the GA of the nine-months-old group at F4, separately for tone and noise segments (Figure 1A and Figure 2A). We chose this procedure because the late negative component was only shown clearly in the responses of the nine-months-old infants. The peak was again set as the most negative point. Amplitudes for the late peak were then measured as the average voltage in 48 ms long windows centered on the GA peak for all infants (see Figure 1A and Figure 2A), separately for responses to tones and noise segments.

Statistical analyses of the difference waveform amplitudes were performed separately for the early and late components and the two stimulus types (tone, noise) by mixed-mode ANOVAs with within-subject factors of ANTERIOR-POSTERIOR (frontal, central, parietal) and LATERALITY (left, central, right), and the between-subject factor of AGE GROUP (four-, nine-months-olds). The scalp distribution factors (ANTERIOR-POSTERIOR and LATERALITY) were based on a matrix of 3 × 3 electrodes: F3, Fz, F4, C3, Cz, C4, P3, Pz, and P4. For post-hoc testing of significant interactions between AGE GROUP and the ANTERIOR-POSTERIOR and/or LATERALITY factor(s) ANOVAs were separately performed for each level of the scalp distribution factors. The false discovery rate (FDR) correction was applied to p-values to account for multiple comparisons ^[51]^. The p values reported reflect the FDR corrected values. The alpha level was set to 0.05. Only significant results are reported. The partial eta squared (ηp^2^) measure of effect size is also reported.

### 4.5. Source localization

The Brainstorm toolbox ^[27]^ was used to perform EEG source reconstruction, following the protocol of previous studies ^[52]^. Age-appropriate templates for both groups were used to derive default anatomical regions ^[25]^, which, along with the default electrode locations, were entered into the forward boundary element head model (BEM) provided by the openMEEG algorithm ^[53]^. The tissue conductivity values were those used in O’Reilly et al. (2021) ^[25]^ in 7-months-old infants (0.33 S/m for the gray matter, 0.0041 S/m for the skull, and 0.33 S/m for the scalp).

For the modeling of time-varying source signals (current density) of all cortical voxels, a minimum norm estimate inverse solution was applied after appropriate noise covariance estimation. Matrices were then scaled through dynamical Statistical Parametric Mapping normalization ^[54]^. Averaging current density across voxels yielded time series for 62 cortical areas (region of interest ROIs), defined by the standardized parcellation scheme introduced by Desikan and Killiany ^[55]^. Finally, for all ROIs, we extracted the average signal for the corresponding ERP peak windows, separately for each SOUND DURATION (short, long), STIMULUS TYPE (tones, noise), and AGE GROUP (4, 9 months).

Based on visual inspection of the source activity on the cortical maps, we selected the bilateral ROIs from the Desikan-Killiany parcellation scheme ^[55]^ for subsequent statistical analysis. <u>Frontal regions:</u> all three anatomical partitions (opercular, orbital, and triangular) of the Inferior Frontal Gyrus (opIFG, orbIFG, triIFG) and the Orbital Frontal Gyrus (OFG) in its medial and lateral parts; <u>parietal regions:</u> the Supramarginal Gyrus (SMG) and the Inferior Parietal Gyrus (IPG); <u>temporal regions:</u> the Superior Temporal Gyrus (STG), the Inferior Temporal Gyrus (ITG), the Middle Temporal Gyrus (MTG), the bank of the Superior Temporal Sulcus (STS), and primary auditory cortex (PAC).

For statistical analyses, average source amplitudes in the selected regions were tested separately for the early and late time windows, for tones and noise segments, and ROIs by between-subjects ANOVAs with the AGE GROUP factor (four-, nine-months-olds). All other aspects of the statistical analysis are identical to that employed for ERPs (see 2.4).

## Authors contributions

Conceptualization, B.T., I.W., and G.H.; methodology, B.T.; formal analysis, S.P.; investigation, G.H.; resources, I.W. and G.H.; data curation, S.P.; writing—original draft preparation, S.P.; writing—review and editing, B.T., I.W., and G.H.; visualization, S.P.; supervision, I.W., H.B.; project administration, B.T., I.W., and G.H.; funding acquisition, I.W. and G.H. All authors have read and agreed to the published version of the manuscript.

## Data Availability Statement

Data available on request due to privacy and ethical restrictions.

