## Supplementary Materials for "Early maturation of sound duration processing in the infant brain"

**Table 1. Testing age effects for source amplitudes in the early time window**
ANOVA results comparing mean CSD values in the time window of the early negative difference, separately for the two stimulus types (tones and noise segments; columns) and ROIs (rows).

| Cortical ROIs | TONES | NOISE SEGMENTS |
|---|---|---|
| **Frontal cortex** | | |
| Right triIFG | - | - |
| Left triIFG | - | F(1,62) = 6; p = 0.017, ηp2 = 0.09 |
| Right opIFG | - | - |
| Left opIFG | F(1,76) = 6.08; p = 0.015, $\eta p^2$ = 0.07 | F(1,62) = 4.91; p = 0.03, ηp2 = 0.07 |
| Right orbIFG | - | - |
| Left orbIFG | - | F(1,62) = 5.07; p = 0.027, ηp2 = 0.08 |
| Right medial OFG | - | - |
| Left medial OFG | - | - |
| Right lateral OFG | - | - |
| Left lateral OFG | - | - |
| **Temporal cortex** | | |
| Right STG | F(1,76) = 4.17; p = 0.044, ηp2 = 0.05 | - |
| Left STG | - | - |
| Right MTG | F(1,76) = 13.06; p = 0.005, ηp2 = 0.15 | - |

| | | |
|---|---|---|
| Left MTG | - | - |
| Right ITG | - | - |
| Left ITG | - | - |
| Right bank of STS | F(1,76) = 9.09; p = 0.003, ηp2 = 0.11 | F(1,62) = 14.57; p = 0.003, ηp2 = 0.19 |
| Left bank of STS | - | - |
| Right PAC | - | - |
| Left PAC | - | - |
| **Parietal cortex** | | |
| Right SMG | - | - |
| Left SMG | - | - |
| Right IPG | - | - |
| Left IPG | - | - |

**Table 2. Testing age effects for source amplitudes in the late time window**
ANOVA results comparing mean CSD values in the time window of the early negative difference, separately for the two stimulus types (tones and noise segments; columns) and ROIs (rows).

| Cortical ROIs | TONES | NOISE SEGMENTS |
|---|---|---|
| **Frontal cortex** | | |
| Right triIFG | - | - |
| Left triIFG | F(1,76) = 8.84; p = 0.003, ηp2 = 0.1 | - |
| Right opIFG | - | - |
| Left opIFG | - | - |
| Right orbIFG | - | - |
| Left orbIFG | | F(1,62) = 4.23; p = 0.043, ηp2 = 0.06 |
| Right medial OFG | - | F(1,62) = 4.86; p = 0.031, ηp2 = 0.07 |
| Left medial OFG | - | - |
| Right lateral OFG | - | - |
| Left lateral OFG | - | F(1,62) = 5.59; p = 0.021, ηp2 = 0.08 |
| **Temporal cortex** | | |
| Right STG | F(1,76) = 11.38; p = 0.0011, ηp2 = 0.13 | F(1,62) = 8.21; p = 0.005, ηp2 = 0.12 |
| Left STG | F(1,76) = 12.44; p = 0.0007, ηp2 = 0.14 | F(1,62) = 5.95; p = 0.017, ηp2 = 0.09 |

| | | |
|---|---|---|
| Right MTG | - | |
| Left MTG | - | F(1,62) = 9.45; p = 0.0031, ηp2 = 0.13 |
| Right ITG | - | - |
| Left ITG | - | F(1,62) = 6.8; p = 0.011, ηp2 = 0.1 |
| Right bank of STS | - | - |
| Left bank of STS | - | F(1,62) = 14.52; p = 0.0003, ηp2 = 0.19 |
| Right PAC | F(1,76) = 12; p = 0.0008, ηp2 = 0.14 | F(1,62) = 12.89; p = 0.0006, ηp2 = 0.17 |
| Left PAC | - | F(1,62) = 6.07; p = 0.016, ηp2 = 0.09 |
| **Parietal cortex** | | |
| Right SMG | F(1,76) = 9.62; p = 0.0026, ηp2 = 0.11 | F(1,62) = 9.58; p = 0.0029, ηp2 = 0.13 |
| Left SMG | - | - |
| Right IPG | - | - |
| Left IPG | - | - |
